# Morphine regulates astrocyte transcriptional dynamics in the ventral tegmental area by stimulation of glucocorticoid signaling

**DOI:** 10.1101/2025.09.22.677814

**Authors:** Jennifer J. Tuscher, Angela Cleere, Robert A. Phillips, Catherine E. Newman, Guy Twa, Nathaniel J. Robinson, Lara Ianov, Robert E. Sorge, Jeremy J. Day

## Abstract

Opioids are potent analgesics often prescribed for the treatment of chronic pain, a condition affecting millions worldwide. Although pain states increase vulnerability to opioid use disorders, the neural mechanisms underlying this interaction remain incompletely understood. The ventral tegmental area (VTA) is a key site for opioid actions, and emerging evidence suggests that pain states and opioid experience both induce transcriptional, molecular, and circuit adaptations in the VTA that contribute to motivated behaviors. However, the transcriptional responses of distinct VTA cell types to each of these factors (alone or in combination) have not been identified. Here, we employed single-nucleus RNA sequencing to comprehensively define transcriptional alterations in the rat VTA to acute morphine administration in a chronic inflammatory pain model. We report that morphine induces gene expression changes primarily in glial cells and dopamine neurons, with minimal effects in other neuronal cell types. Surprisingly, VTA astrocytes and oligodendrocytes exhibited the most robust transcriptional responses to opioid exposure, despite lacking detectable opioid receptor expression. Among the most highly regulated glial genes was *Fkbp5*, which encodes a co-chaperone protein that acts in concert with heat shock proteins to modulate stress responses. Using pharmacological and CRISPR-based approaches in rat glial cells and human astrocytes, we demonstrate that regulation of *Fkbp5* is mediated indirectly through glucocorticoid signaling rather than direct opioid receptor activation. These findings reveal that glial cells within reward circuits undergo profound transcriptional reprogramming in response to opioids through indirect, stress-hormone mediated mechanisms, highlighting a previously unappreciated non-neuronal contribution to opioid-induced neural adaptations.

## INTRODUCTION

Opioid use disorder (OUD) represents a pressing public health challenge, with overdose deaths attributed to opioids reaching unprecedented levels in the last decade. While the acute reinforcing effects of opioids are well-established, the molecular mechanisms underlying the transition from initial use to dependence remain incompletely understood. The ventral tegmental area (VTA), a critical component of the mesolimbic reward circuit, has long been recognized as a key brain region mediating opioid reinforcement and addiction-related behaviors^1–4^. Within the VTA, opioids are thought to act at μ-opioid receptors (μORs) expressed in local GABAergic interneurons, causing a disinhibition of dopamine neurons^5–7^. Additionally, chronic pain induces adaptations in VTA dopamine neurons and glutamatergic inputs to dopamine neurons, which contribute to the development of anhedonia-like behaviors observed in pain states^8,9^. However, emerging evidence suggests that non-neuronal cell populations, particularly glia, may play equally important roles in mediating the long-term consequences of opioid exposure and the pathophysiology of chronic pain conditions^10–24^.

Glial cells, including astrocytes, microglia, and oligodendrocytes, comprise the majority of cells in the central nervous system and are increasingly recognized as active participants in synaptic transmission^11,12,25–27^, neuroplasticity^13,15,28–30^, and drug-induced behavioral adaptations^10,13,14,27,30,31^. Recent transcriptomic studies in both preclinical models^17,22,23^ and human postmortem tissue^18–21^ have revealed robust opioid-induced gene expression changes in glial populations, further suggesting these cells may contribute significantly to the molecular pathology underlying OUD. Of particular interest is the emerging role of stress-responsive pathways in mediating opioid-induced transcriptional changes. Notably, the hypothalamic-pituitary-adrenal (HPA) axis and glucocorticoid signaling pathways have been implicated in addiction vulnerability^32,33^, with mounting evidence suggesting that opioid exposure can engage stress response machinery even in the absence of external stressors^34,35^. Further, chronic pain conditions, which often precede the development of OUD in clinical populations, may prime the brain for enhanced sensitivity to opioid-induced molecular changes.

The interaction between pain-related neuroadaptations and subsequent opioid exposure remains poorly characterized at the cellular and molecular level, particularly with respect to cell-type-specific responses within reward-related brain regions. To address these knowledge gaps, we employed single-nucleus RNA sequencing (snRNA-seq) to generate a comprehensive molecular atlas of cell-type-specific transcriptional responses to chronic pain and acute morphine exposure in the rat VTA. This high-resolution approach provided unprecedented insight into the heterogeneous responses of different VTA cell populations to these treatments. Our findings reveal that glial cells, rather than neurons, exhibit robust transcriptional responses to opioid exposure, with particular enrichment of heat shock and stress-related gene signatures. Through complementary *in vitro* experiments using novel CRISPR interference (CRISPRi) tools for cell-type-specific gene perturbation, we provide mechanistic insights into the glucocorticoid-mediated pathways that drive these glial responses and their potential relevance to OUD.

## RESULTS

### Single nucleus RNA profiling reveals cell-type-specific transcriptional responses to pain and opioid experience in the rat VTA

To examine how chronic pain and acute morphine impact cell-type-specific transcriptional responses in the rat VTA, we used the 10x Genomics Chromium platform to perform snRNA-seq on 50,513 nuclei collected from 14 male and 14 female Sprague-Dawley rats (mean = 3,608 nuclei per GEM well). Rats were first injected with either vehicle or Complete Freund’s Adjuvant (CFA; 100% i.p.l.) into the hind paw, resulting in chronic inflammation that leads to thermal and mechanical hyperalgesia^36,37^. On day 7, vehicle and CFA groups were then treated with either saline or morphine (10 mg/kg), yielding 4 treatment groups: 1) vehicle + saline, 2) vehicle + morphine, 3) CFA + saline, 4) CFA + morphine (**Fig. 1a-b**). Pain sensitivity was measured at baseline and on days 2 and 7 of the experimental timeline using the Von Frey test. Prior to CFA injection, all rats exhibited similar withdrawal thresholds at baseline (**Fig. 1c**). However, by 24 hours after injection, CFA-treated rats showed a significant reduction in withdrawal threshold relative to vehicle-treated rats, indicative of elevated pain sensitivity. This heightened pain state was maintained at day 7, the final measurement recorded prior to treatment with saline or morphine. One hour after saline or morphine injections, tissue was collected for snRNA-seq to examine experience-dependent transcriptional changes in the rat VTA. Dimensionality reduction and clustering of the final integrated object containing 38,468 nuclei resulted in 16 transcriptionally distinct populations (**Fig. 1d**), including dopaminergic (DA.1 and DA.2), glutamatergic (Glut.1, Glut.2, and Glut.3), and GABAergic (GABA.1 and GABA.2) neuronal populations previously identified in the rat VTA^38^ (see methods for full details regarding dimensionality reduction parameters, doublet filtering, ambient RNA removal, quality filtering, and removal of nuclei from adjacent brain regions). Quality control, integration, and marker gene expression within defined clusters can be found in **Fig. S1**. Among both neuronal and non-neuronal VTA cell classes, cell-type annotations align with much larger recently published datasets from the mouse brain^39^ (**Fig. S2**), suggesting transcriptional conservation of VTA cell identities across rodent species. Consistent with our prior work examining the rat VTA^38^, we also observed two transcriptionally distinct dopaminergic subpopulations: DA.1, which exhibits preferential enrichment of *Gch1*, a marker of canonical DA neurons, and DA.2, a combinatorial population enriched with *Slc26a7* and molecular markers of glutamatergic and GABAergic machinery (**Fig. S3**).

**Figure 1.**
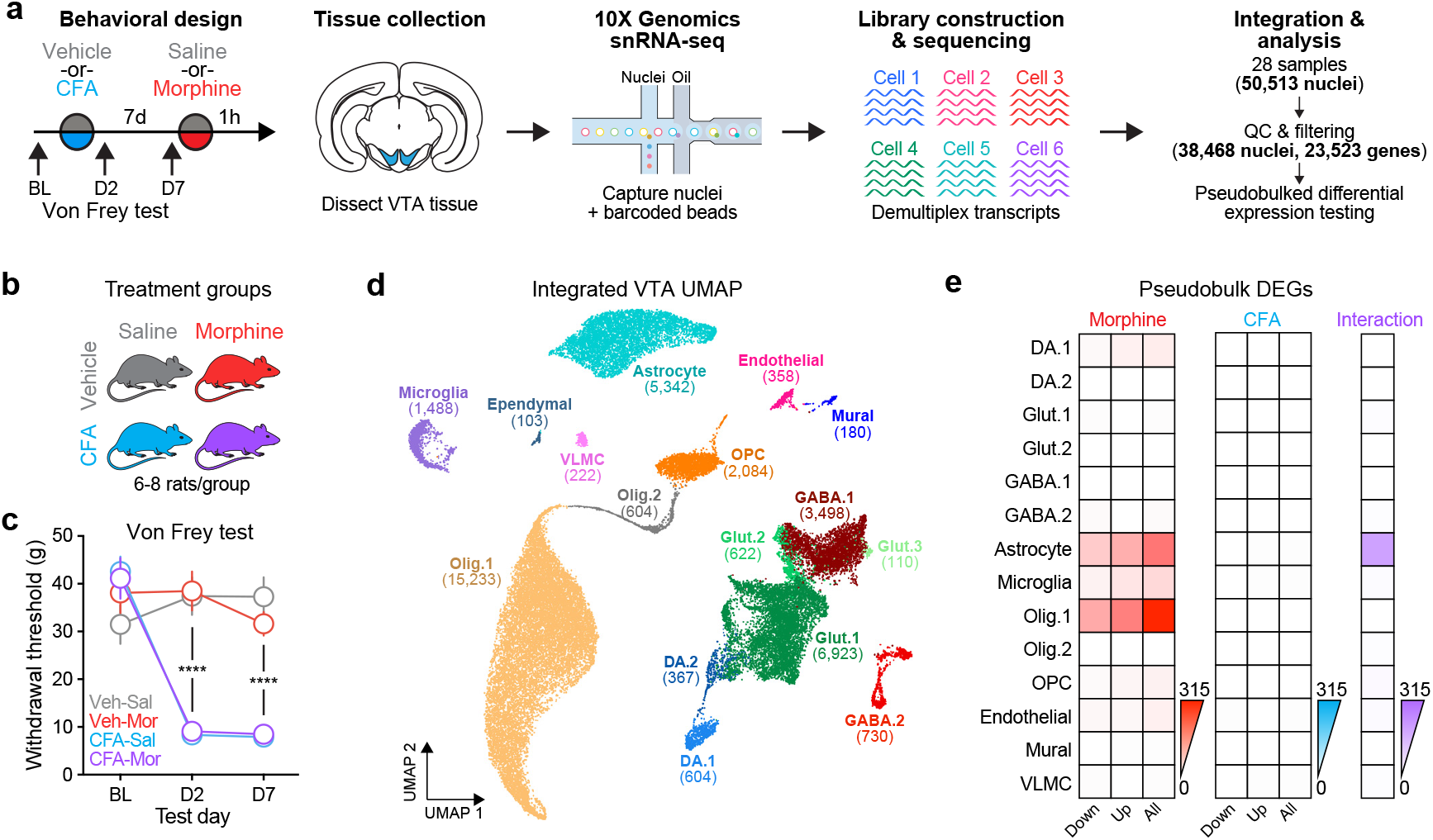
Morphine induces gene expression changes in specific VTA cell populations. **a**, Illustration of behavioral and snRNA-seq workflow. **b**, Treatment groups and sample sizes for snRNA-seq. **c**, Behavioral effects of CFA-induced inflammatory pain measured with the Von Frey test. CFA induced mechanical hyperalgesia on day 2 and day 7 after administration (two way ANOVA, significant interaction between test day and CFA treatment, F(6, 108)=14.61; p<0.0001, with Tukey post-hoc tests revealing group differences on D2 and D7). **d**, Integrated VTA snRNA-seq UMAP from 28 independent samples, totalling 38,468 nuclei after quality control and filtering. Clustering identified 16 distinct cell types. **e**, Total number of differentially expressed genes (DEGs) identified using cluster-specific pseudobulked analysis with DESeq2.

After initial validation of global clustering using cell-type-defining gene markers (**Fig. S1e**), differentially expressed gene (DEG) analysis for main treatment effects and treatment interactions (with sex as a covariate) was completed using pseudobulked count matrices in DESeq2 v.1.44.0^40^ (likelihood ratio test, adjusted *p* value < 0.05). Overall, transcriptional responses to persistent pain induced by CFA were limited, with only two DEGs passing statistical significance criteria (**Fig. 1e, Table S1**). In contrast, acute morphine treatment resulted in robust transcriptional alterations across several VTA cell populations, but were most notable in DA.1 (27 DEGs), Astrocyte (198 DEGs), Olig.1 (313 DEGs), and Microglia (57 DEGs) clusters (**Fig. 1e, Table S1**). Interactions between morphine and chronic pain state were most frequently observed in the astrocyte cluster, where 201 DEGs were identified. Overall, these findings reveal diverse transcriptional responses to acute morphine and chronic pain experience across the various cell populations that comprise the rat VTA, with glial populations exhibiting the largest number of changes.

### Morphine induces experience-dependent gene expression changes in dopamine neurons

The primary receptor target of abused opioids such as morphine, fentanyl, heroin, and oxycodone is the mu opioid receptor (µOR), which is encoded by the *Oprm1* gene. Deletion of *Oprm1* abolishes the rewarding, analgesic, and withdrawal-related effects of opioids^41,42^, and the VTA has long been considered a target for the reinforcing properties of opioids^2,5,6,43–46^. However, while µOR activity has traditionally been linked to suppression of GABAergic interneurons in the midbrain^5,47^, few studies have comprehensively characterized *Oprm1* expression across molecularly defined VTA neuronal subtypes. Therefore, we first sought to examine expression patterns of *Oprm1* across dopaminergic, glutamatergic, and GABAergic VTA neurons (**Fig. 2a-c**). In agreement with prior molecular characterization studies, *Oprm1* mRNA was nearly absent in dopamine neurons, and was expressed at low levels in several glutamatergic and GABAergic clusters^38,47^. We observed abundant *Oprm1* mRNA in the GABA.2 neuronal population, which also expressed well-defined markers of GABAergic interneurons such as *Nos1, Kit, Sst*, and *Vip* (**Fig. S4a-c**). Within this neuronal cluster, *Oprm1* was co-expressed in over 80% of the nuclei that also expressed interneuron markers *Reln, Nts, Calb1, Kit*, and *Sst*, and was rarely co-expressed in *Vip*+ neurons (**Fig. S4d**). Although many of these genes have been referred to as interneuron markers in the literature^5,7,48^, more recent evidence suggests these populations may instead represent GABAergic neurons that make long-range projections in addition to intra-VTA synaptic connections^49^. Additionally, it is possible that the GABA.2 neuronal population overlaps with previously described GABAergic inputs from the “tail” of the VTA, also called the rostrotegmental nucleus (RMTg), which exhibits high expression of µORs and projects to VTA dopamine neurons^45,50,51^. Nevertheless, these results highlight that mRNA for the µOR is enriched in a specific GABAergic neuronal subpopulation, and is not found in dopamine neurons.

**Figure 2.**
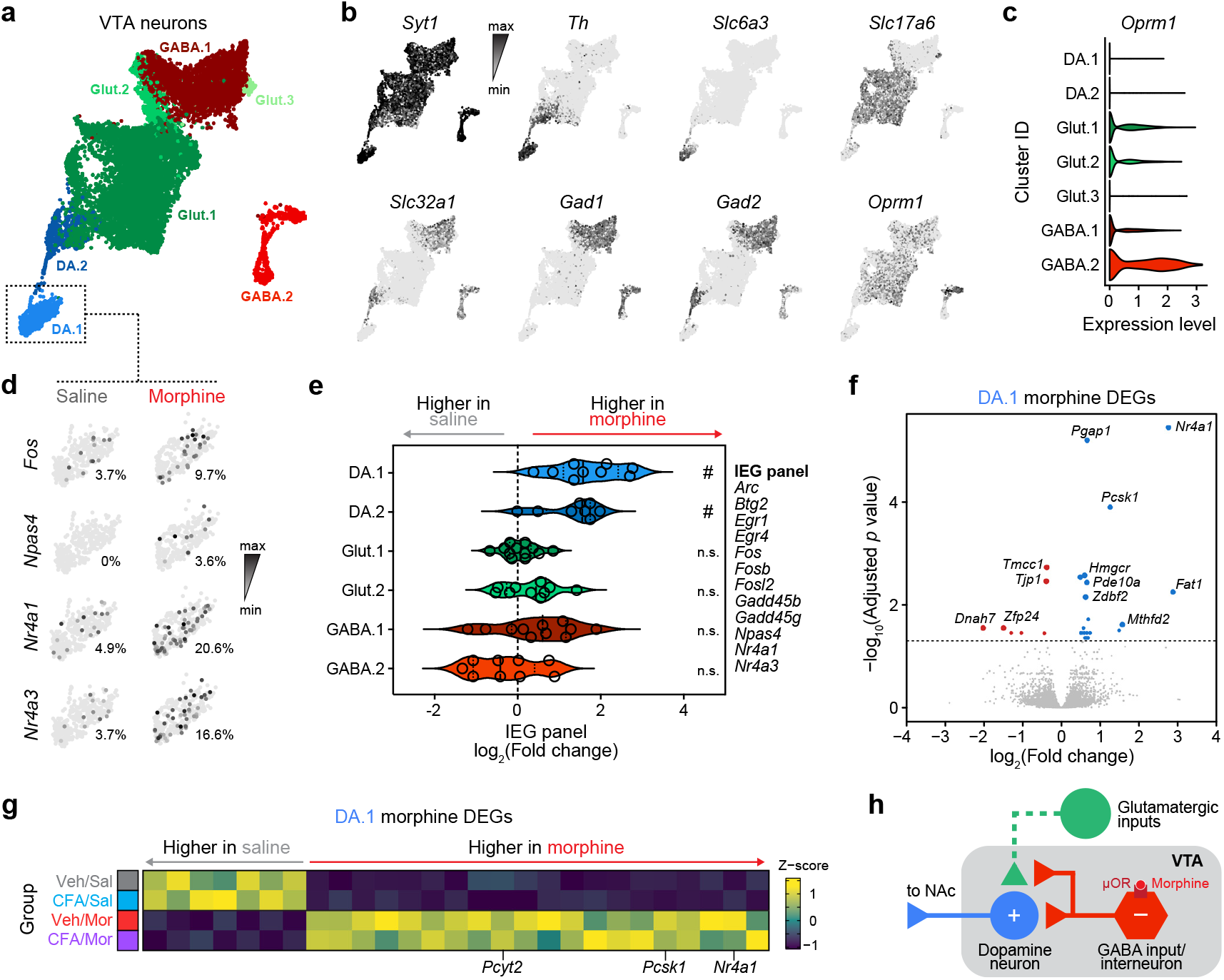
Morphine induces immediate early genes in dopamine neurons. **a**, UMAP representation of neuronal cluster IDs. **b**, Marker genes for neurons (*Syt1*) dopamine neurons (*Th, Slc6a3*), glutamate neurons (*Slc17a6*), and GABA neurons (*Slc32a1, Gad1, Gad2*), as well as the µ opioid receptor gene *Oprm1*, plotted with UMAP coordinates (from **a**). **c**, Violin plot of expression for *Oprm1* across neuronal cell types reveals highest expression in GABA.2 neurons. **d**, Expression of immediate early genes in DA.1 neuronal cluster, split by saline or morphine treatment. **e**, Expression of IEG module genes across VTA cluster IDs reveals selective upregulation in dopamine neurons. **f**, Volcano plot of DA.1 morphine DEGs identified from pseudobulk analysis with DESeq2. **g**, Heatmap of DA.1 morphine DEGs across all experimental groups reveals no systematic interaction between pain state and morphine transcriptional response. **h**, Illustration of circuit model for dopamine neuron activation by morphine.

Although VTA dopamine neurons do not express *Oprm1* (**Fig. 2b-c**), a classic model of opioid action posits that VTA dopamine neuron activity is likely modulated by opioids via disinhibition that results from silencing of GABAergic interneurons and/or reduced inhibition of excitatory synapses onto dopamine neurons^52,53^. To explore this hypothesis, we examined the levels of known immediate early genes (IEGs) in dopamine neurons between saline and morphine administration groups. As expected, we observed that an elevated percentage of neurons in the DA.1 population expressed IEGs such as *Fos, Npas4, Nr4a1*, and *Nr4a3* following morphine administration (**Fig. 2d**). Strikingly, morphine increased expression of a panel of IEGs in both DA.1 and DA.2 populations (**Fig. 2e**), which differ in the relative expression of dopamine, glutamatergic, and GABAergic markers as previously reported^38^ (**Fig. S3**). We found no change in IEG expression in other neuronal populations (**Fig. 2e**). Known activity-responsive genes such as *Nr4a1*^*54,55*^ and *Pcsk1*^*56*^ also passed stringent statistical cutoffs to qualify as DEGs in dopamine neurons (**Fig. 2f**), although other IEGs (*Fos, Npas4*, and *Nr4a3*) missed statistical thresholds for significance, likely due to dropout or expression in a low overall percentage of dopamine neurons. We observed no interaction between pain state and morphine administration in DEGs identified in the DA.1 neuronal cluster (**Fig. 2g**). Overall, these results are consistent with a model in which opioids disinhibit dopamine neurons via actions at GABAergic inputs or GABAergic interneurons in the VTA (**Fig. 2h**).

### VTA glial populations exhibit robust transcriptional responses to opioid experience

Other than microglia, VTA glial cell populations do not exhibit robust expression of any genes encoding opioid receptors (**Fig. 3a**). Nevertheless, pseudobulked DEG analysis revealed robust changes in gene expression in several non-neuronal cell populations, led by the Astrocyte, Olig.1, and Microglia clusters (**Fig. 1e, Table S1**). Genes such as FK506 binding protein 5 (*Fkbp5*), a prolyl isomerase in the immunophilin gene family, were strongly induced by morphine administration in all 3 of these glial populations (**Fig. 3b-c**). Further analysis suggested shared induction of upregulated DEGs and minimal overlap of downregulated DEGs (**Fig. 3d**). In addition to *Fkbp5*, shared morphine-induced genes included *Bag3, Dnaja4*, and *Hsph1*, all of which are involved in protein folding and chaperone-binding activity. Gene ontology analysis conducted with gene sets induced by morphine revealed an enrichment in processes related to protein modification, localization, and folding, as well as heatshock protein binding (**Table S2**). Downregulated genes were enriched for biological categories related to axon guidance, regulation of actin cytoskeleton organization, and Wnt signaling pathways. Despite recent reports of robust hormone-driven sex differences in pain and opioid responses^57^, we observed largely similar morphine-induced gene changes in both male and female samples (**Fig. S5**). To formalize sex comparisons, we repeated DESeq2 analysis with sex as a factor (instead of a covariate), and observed no sex x treatment interactions in any glial cell population. Only one sex x treatment interaction was identified in the DA.2 neuronal population, with no other significant DEGs detected across all other cell populations.

**Figure 3.**
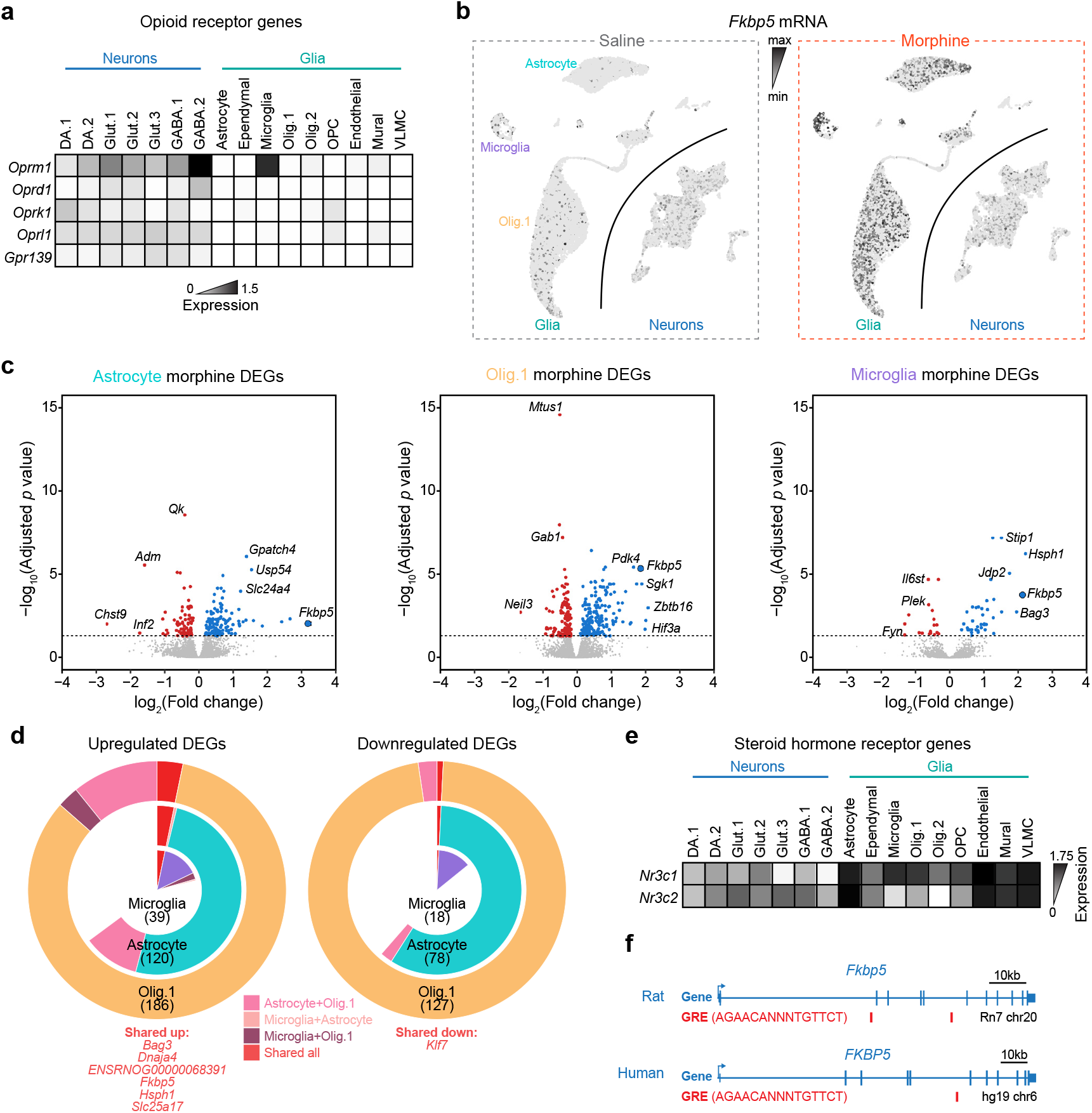
Despite low or absent expression of opioid receptors, morphine strongly alters gene expression in VTA non-neuronal cell populations. **a**, Expression heatmap for opioid receptor genes across VTA cluster IDs. **b**, Feature plot of *Fkbp5*, split by experimental group (saline and morphine). **c**, Volcano plots for differentially expressed genes in astrocytes, oligodendrocytes, and microglia. **d**, Pie charts showing the intersection of upregulated and downregulated DEG lists across three glial populations. e, Expression heatmap for steroid hormone receptor genes *Nr3c1* and *Nr3c2* across VTA cluster IDs. **f**, Gene track for shared DEG *Fkbp5* identifies glucocorticoid response element (GRE) in both the rat (top) and human (bottom) genome.

While morphine-responsive DEGs were distributed across glial populations, interactions between morphine and pain state were relatively concentrated in the Astrocyte cluster (201 interaction DEGs versus 28 interaction DEGs in all other clusters combined; **Table S1**). Interaction DEGs exhibited two general patterns (**Fig. S6a-b**). Group A genes were suppressed by the chronic pain state in drug-free conditions, and this suppression was relieved by morphine administration. Group B genes were increased by CFA-induced chronic pain, and this increase was similarly normalized by morphine administration. Whereas Group A genes were enriched for gene ontology pathways related to mitogen activated protein kinase signaling, Group B genes were enriched in two distinct biological pathways centered on 1) mitochondrial function and the electron transport chain, and 2) ribosomal function (**Fig. S6c, Table S2**). These results suggest that CFA-induced inflammatory pain may generate an astrocyte state that requires increased metabolic demand and de novo protein synthesis, and that this state is (at least temporarily) reversed by opioid administration.

The observation that the most robust morphine-induced gene expression changes were observed in astrocyte and oligodendrocyte populations that lack mRNA for opioid receptor genes (**Fig. 3a**) suggests that these transcriptional alterations are mediated via an indirect mechanism of action. Previous work has identified a role for glucocorticoid receptors (GRs) in mediating transcriptional changes at genes such as *Fkbp5*^*58*^, and this relationship has also been implicated in psychiatric disorders^59^. Given that morphine administration elevates corticosterone levels on an acute timescale^34^, we hypothesized that morphine-induced activation of the hypothalamic-pituitary-adrenal (HPA) axis may drive the observed glial transcription responses through glucocorticoid signaling. To investigate this possibility, we next examined glucocorticoid and mineralocorticoid receptor expression across all VTA populations, and found *Nr3c1* and *Nr3c2* were highly expressed in glial populations (**Fig. 3e**). Further supporting this putative mechanism, glucocorticoid response elements (GREs) have previously been identified within human and rodent *Fkbp5* (**Fig. 3f**), suggesting a potential direct mode of regulation at this locus^60–63^.

### Fkbp5 induction is mediated by glucocorticoid signaling and not µOR activation in rat glia

To test this hypothesis, we first used a C6 rat glial cell culture model system to ask whether the transcriptional responses observed *in vivo* could be recapitulated via activation of glucocorticoid signaling *in vitro*. We used *Fkbp5* as a transcriptional read out, as this gene yielded one of the largest changes in expression after morphine exposure in astrocytes, microglia, and oligodendrocytes in the rat VTA, and has previously been linked to GR signaling^60^. We found dose-dependent increases in *Fkbp5* expression in C6 cells after treatment with corticosterone (**Fig. 4a**), which were blocked by a 30 minute pre-treatment with the GR antagonist mifepristone (**Fig. 4b**). These transcriptional responses could not be induced with the selective µOR agonist DAMGO (**Fig. 4c**). Collectively, these findings demonstrate that glucocorticoid signaling, rather µOR opioid receptor activation, is sufficient for the induction of heat shock associated gene *Fkbp5* in rat glial populations.

**Figure 4.**
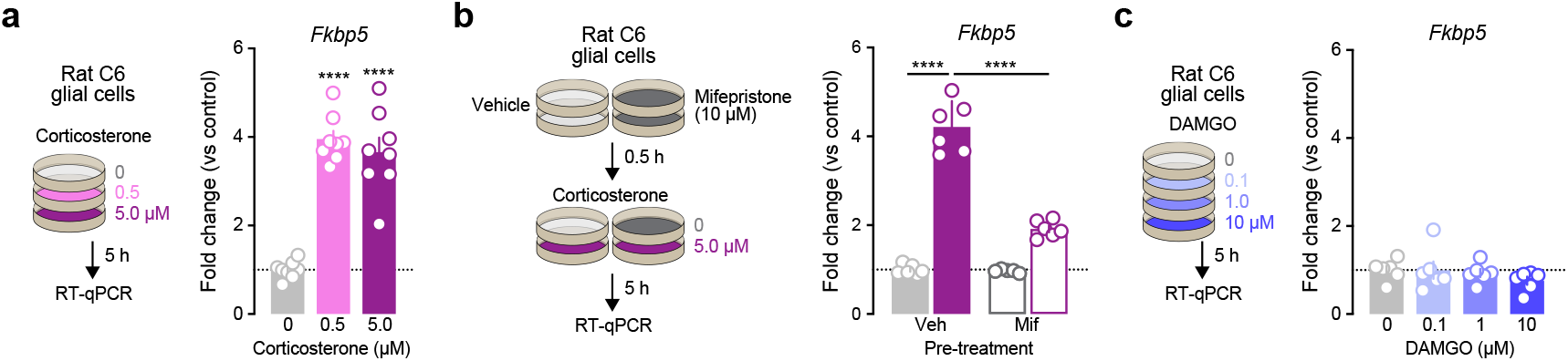
Corticosterone (but not µOR activation) induces *Fkbp5* in rat glial cells. **a**, Treatment with corticosterone, an endogenous glucocorticoid in rats, increases *Fkbp5* expression as measured by RT-qPCR in rat C6 glioma cells (One-way ANOVA, n = 8 per group). **b**, Pretreatment with the glucocorticoid receptor antagonist mifepristone 0.5 h prior to application of corticosterone attenuated the induction of *Fkbp5* (One-way ANOVA, n = 5-6 per group). **c**, Treatment with the µOR agonist DAMGO for 5 h failed to induce *Fkbp5* (One-way ANOVA, n = 6 per group). Data expressed as mean ± s.e.m. Multiple comparisons, ****p < 0.0001.

### FKBP5 is regulated by glucocorticoid signaling in human astrocytes

To expand the translational relevance of our findings, we developed a human-derived astrocyte cell culture model system to dissect the potential role of glucocorticoid regulation of *FKBP5*. Neural precursor cells derived from the human ventral midbrain were differentiated into astrocyte-like cells that expressed GFAP and S100B (two commonly used markers of astrocytes; **Fig. 5 a-b**). Again using *FKBP5* as a transcriptional read out, we treated differentiated human astrocytes with cortisol or the glucocorticoid agonist dexamethasone *in vitro*. Consistent with our observations in rat C6 cells, we found that both cortisol (**Fig. 5c**) and dexamethasone (**Fig. 5d**) were sufficient for *FKBP5* induction in human astrocytes. Agonism of µORs with DAMGO was similarly unable to induce *FKBP5* in our human culture system (**Fig. 5e**). Finally, mifepristone pre-treatment experimental results mirrored observations in rat C6 cells, blocking *FKBP5* upregulation typically observed after cortisol (**Fig. 5g**) or GR activation with dexamethasone (**Fig. 5h**).

**Figure 5.**
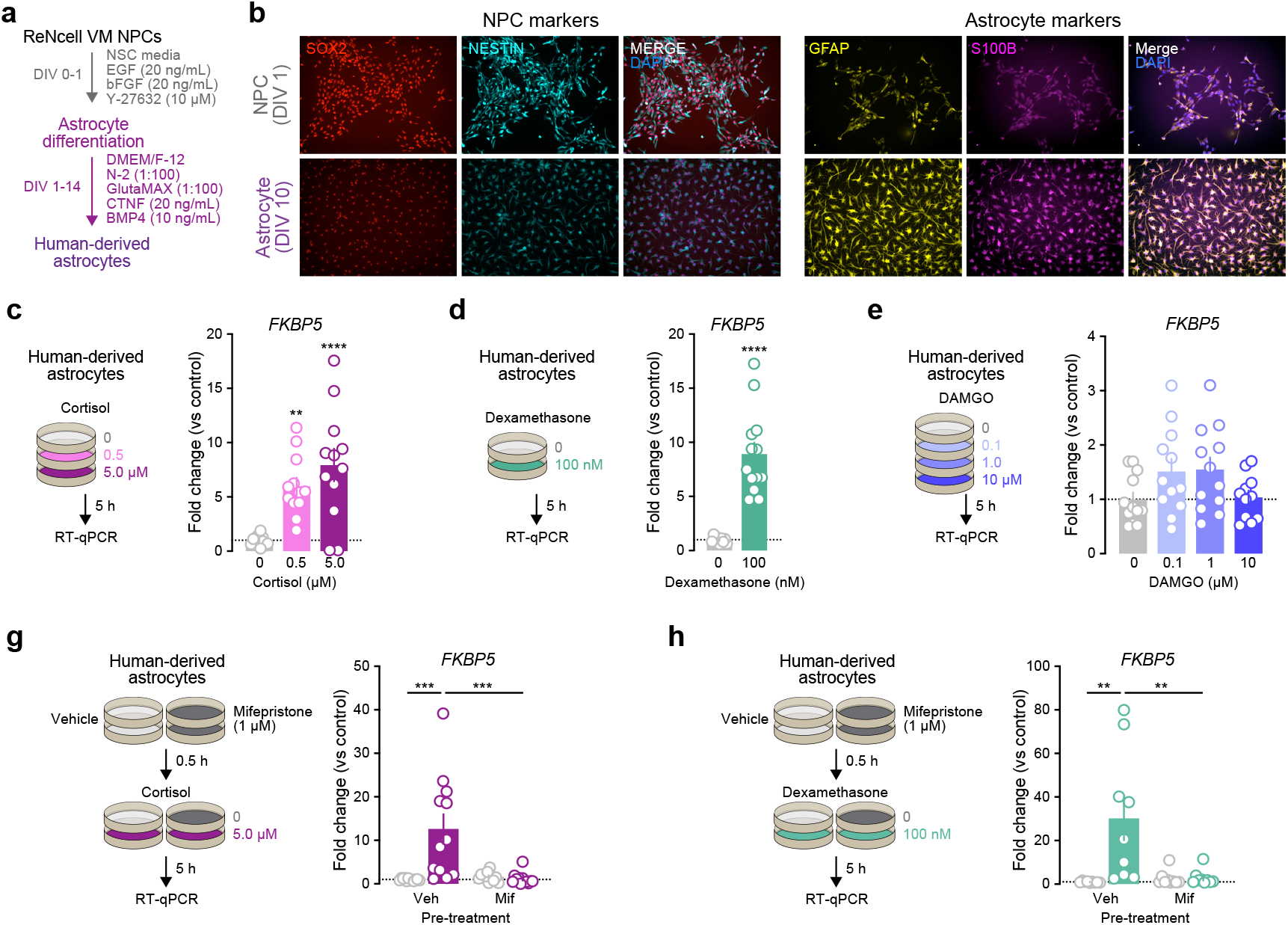
Cortisol (but not µOR activation) induces *FKBP5* in a human-derived astrocyte model via glucocorticoid receptor signaling. **a**, Illustration for generation of human-derived astrocytes from immortalized neural precursor cells (NPCs). **b**, Immunocytochemistry images showing acquisition of astrocyte markers GFAP and S100B across the differentiation time course, as well as depletion of NPC markers. **c**, Treatment with cortisol increases *FKBP5* expression as measured by RT-qPCR in human-derived astrocytes in a dose-dependent manner (One-way ANOVA, n = 12 per group). **d**, Treatment with the glucocorticoid receptor agonist dexamethasone induces *FKBP5* expression as measured by RT-qPCR in human-derived astrocytes (Unpaired t-test, n = 13 per group). **e**, Treatment with the µOR agonist DAMGO for 5 h failed to induce *FKBP5* (One-way ANOVA, n = 15 per group). **f**, Pretreatment with the glucocorticoid receptor antagonist mifepristone 0.5 h prior to application of cortisol blocked the induction of *FKBP5* (One-way ANOVA, n = 12 per group). **h**, Mifepristone blocked dexamethasone-induced increases in *FKBP5* in human astrocytes. Data expressed as mean ± s.e.m. Multiple comparisons, *p < 0.05, **p < 0.01, ***p < 0.001, ****p < 0.0001.

One caveat to using the GR antagonist mifepristone is that it lacks both receptor and cell-type specificity, in that it can have off-target effects on progesterone receptors and is not restricted to astrocytes. To overcome this issue, we developed a lentiviral version of a CRISPR interference (CRISPRi) system that relies on a truncated GFAP promoter for targeted gene repression in astrocytes (**Fig. 6a**). To selectively knock down GR expression in human astrocytes, we first designed and validated single guide RNAs (sgRNAs) that target the human sequence for *NR3C1*. To confirm successful repression using astrocyte selective CRISPRi, human-derived astrocytes were co-transduced with lentiviral GfaABC1D-FLAG-dCas9-KRAB-MeCP2 and sgRNAs targeting either human *NR3C1* or the non-targeting control *lacZ*, yielding a ∼55% reduction in *NR3C1* (**Fig. 6b-c**). We next repeated this approach, followed by treatment with cortisol on DIV14. Here, we found that selective *NR3C1* repression in human astrocytes ablated cortisol-mediated induction of *FKBP5* (**Fig. 6d**). Collectively, our results reveal that transcriptional alterations observed in astrocytes after morphine exposure are likely mediated indirectly via glucocorticoid signaling rather than directly through µORs, and that this mechanism is conserved in human astrocytes.

**Figure 6.**
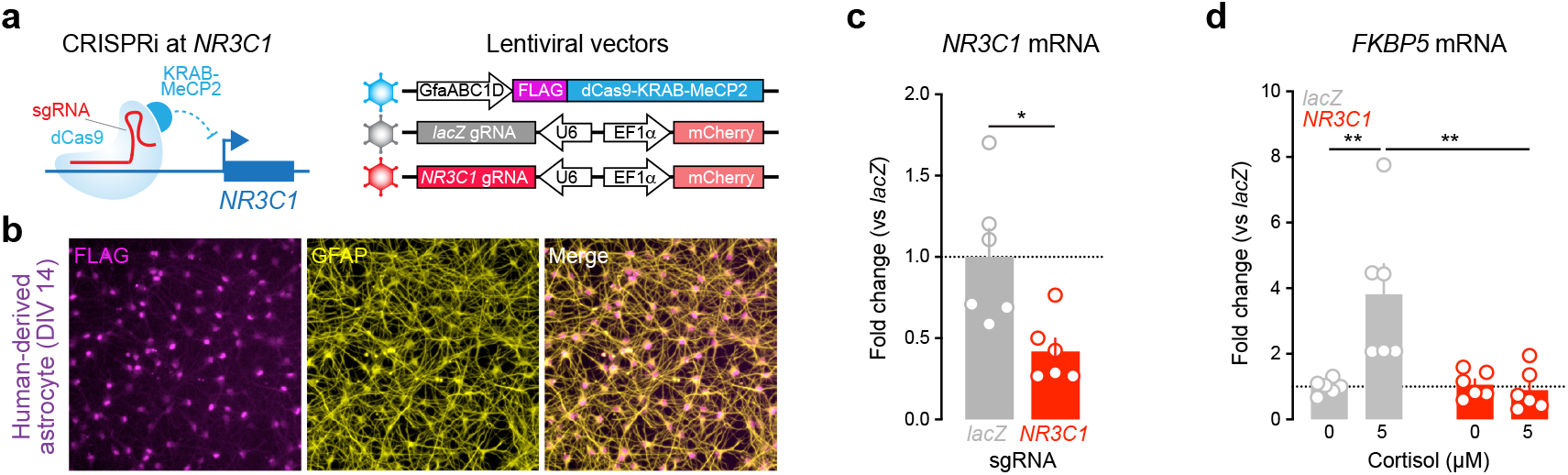
Cortisol induction of *FKBP5* in a human-derived astrocyte model requires *NR3C1*. **a**, Illustration for CRISPRi targeting the *NR3C1* promoter. Lentiviral vectors express a dCas9-KRAB-MeCP2 fusion from the astrocyte-selective GfaABC1D promoter, and CRISPR sgRNAs for *lacZ* (a non-targeting control) and *NR3C1* are delivered via separate viral vectors. **b**, Immunocytochemistry images showing expression of CRISPRi machinery in human-derived astrocytes. **c**, *NR3C1* sgRNA paired with CRISPRi machinery successfuly reduces *NR3C1* mRNA as compared to *lacZ* control sgRNA (Mann-Whitney nonparametric test,, n = 6 per group). **d**, *NR3C1* CRISPRi blocked the induction of *FKBP5* by cortisol in human-derived astrocytes (Two-way ANOVA, n = 6 per group). Data expressed as mean ± s.e.m. Multiple comparisons, *p < 0.05, **p < 0.01.

## DISCUSSION

Opioid use disorder remains a significant public health crisis both in the United States and globally, with over 100,000 opioid-related overdose fatalities reported in the U.S. in recent years (CDC). Many individuals who develop OUD are initially prescribed opioids for chronic pain or other medical conditions, but go on to develop a dependence due to the addictive nature of these substances. Despite the availability of life-saving interventions including methadone, naltrexone, and buprenorphine, many individuals continue to struggle with OUD, highlighting the continued need for innovative therapeutic approaches that target the underlying biological mechanisms engaged by opioids which may ultimately lead to dependence.

Here, we provide a high-resolution molecular atlas for cell-type specific transcriptional responses to pain and morphine in the rat VTA. Using snRNA-seq, we explored over 50,000 nuclei from male and female Sprague-Dawley rats subjected to chronic pain and/ or acute morphine treatment conditions. Our findings reveal distinct and heterogeneous transcriptional responses across various cell populations in the VTA, highlighting the complex molecular adaptations to pain and opioid exposure, particularly in glial clusters. Although limited in comparison to glial populations, we also observed a small number of IEGs engaged in DA neurons in response to acute morphine, consistent with a disinhibition mode of action. To expand on our sequencing observations, we next examined the mechanisms underlying heat shock-related gene expression in both rodent and human-derived glial culture systems. Our results are consistent with a role for glucocorticoid signaling in glial cell types rather than direct regulation by µORs. Finally, to refine cell type and receptor selectivity, we used a novel CRISPRi approach for transcriptional perturbation in astrocytes. Collectively, these findings add to the growing literature implicating glial populations in mediating molecular adaptations to pain and opioid use within mesolimbic reward circuitry.

Interestingly, transcriptional changes induced by chronic pain were exceptionally limited across all cell populations, with only 2 DEGs surviving statistical cutoffs. This lack of transcriptomic response may be reflective of the delayed time point (7 days after CFA injection) assayed for pain conditions, rather than an indication that CFA-induced pain states yield no impact on gene expression in the VTA. This notion is consistent with prior observations of IEG induction in the VTA on an acute time frame following exposure to other forms of painful stimuli^64–66^. In contrast, acute morphine treatment elicited widespread transcriptional alterations in glial populations, with shared engagement of heat shock associated gene signatures between astrocytes, microglia, and oligodendrocytes. These findings underscore the robust impact of opioids on glial cell function and their potential role in modulating reward neurocircuitry in the context of drug exposure. Our observations align with recent studies also highlighting robust opioid-induced gene changes in glial populations within rodent model systems^17,22,23,67,68^ and human opioid users^19–21^.

Despite limited overall neuronal transcriptomic responses to prolonged pain or acute morphine, dopamine neurons exhibited modest changes in several immediate early genes (e.g., *Fos, Npas4, Nr4a1*, and *Nr4a3)* following morphine exposure. The absence of *Oprm1* expression in dopamine neurons suggests these changes are mediated via an indirect mechanism, likely involving disinhibition of locally synapsing GABAergic neurons. Enrichment of *Oprm1* in GABAergic neuron clusters supports this hypothesis, and is consistent with prior reports of dopaminergic disinhibition in the VTA^5,69^.

Notably, the vast majority of transcriptomic changes in our dataset, regardless of treatment group, occurred within glial populations. Further analysis of morphine-induced DEGs revealed enrichment of genes related to heat shock signaling across major glial clusters including astrocytes, microglia and oligodendrocytes. While microglia exhibit moderate levels of *Oprm1*, similar levels of expression were not observed in astrocytes or oligodendrocytes (**Fig. 3a**), populations which exhibited the largest number of significant transcriptional changes overall. Surprisingly, effects on gene expression in these glial populations appear to be mediated via GRs rather than direct engagement of µORs. This notion is corroborated by *in vitro* experiments using rat glial and human-derived astrocyte culture systems, where engagement of glucocorticoid signaling, rather than µOR agonism, induced the expression of *FKPB5*, a key gene implicated in this pathway.

In recent years, the role of *Fkbp5* in modulating stress responses and addiction-like behaviors has garnered significant attention. As a key negative regulator of glucocorticoid signaling, *Fkbp5* has important implications for regulation of the stress response and the etiology of several neuropsychiatric conditions, including SUDs. Genetic variations in the human *FKBP5* gene are associated with individual susceptibility to addiction, particularly in the context of opioid use. For example, 2 intronic *FKBP5* single nucleotide polymorphisms (SNPs) have been linked to long-term heroin use in a clinical cohort of over 1,000 subjects with European or Middle-Eastern ancestry, an observation replicated in a smaller independent African American cohort^70^. Moreover, recent single-cell profiling work from human opioid users has also observed upregulation of *FKBP5* in astrocytes, oligodendrocytes and oligodendrocyte precursor cells, particularly in women with OUD^19^. This further reinforces the relevance of *FKBP5* in the context of OUD and suggests it may serve as an important target for therapeutic intervention.

Preclinical models complement these human findings, demonstrating consistent *Fkbp5* dysregulation across several brain regions and opioid exposure paradigms. For example, *Fkbp5* expression is upregulated in the mouse striatum after both acute^67,68^ and chronic opioid administration^71^ and in response to oxycodone^72^. Similarly, chronic morphine and precipitated withdrawal both increase *Fkbp5* expression in the locus coeruleus and VTA^67,73^, and suppression of FKBP5 was found to attenuate the development of dependence. In line with these observations, recent single-cell profiling efforts targeting the rat nucleus accumbens have also identified increases in *Fkbp5*^*20,22*^ in glial populations. Further, these results dovetail with recent rodent findings demonstrating that treatment with the glucocorticoid receptor antagonist mifepristone (which blocked *Fkbp5* induction in our cell culture studies) also significantly reduced heroin consumption and motivation for heroin in an addiction-relevant intravenous self-administration model^74^. Taken together, these findings highlight a central role for *Fkbp5* in modulating biological pathways that underlie both stress and addiction, making it a key target for therapeutic interventions.

To further examine the requirement of GR activation in driving the expression of *Fkbp5* in astrocytes, we developed a CRISPRi system for selective perturbation of this population. This construct relies on the GfaABC1D promoter, a truncated version of the *Gfap* promoter which allows for improved selectivity in targeting astrocytes^75^. Our observations in human astrocytes highlight the translational relevance of glucocorticoid signaling in opioid responses, suggesting potential therapeutic targets for modulating glial responses to opioid exposure. The development and application of this CRISPRi approach to selectively knockdown GR expression in astrocytes further supports the specificity of glucocorticoid-mediated transcriptional regulation in these cells.

In summary, our results allow three broad conclusions. First, opioid exposure produces distinct and mechanistically independent transcriptional changes in dopamine neurons and glia resident to the VTA. In dopamine neurons, morphine induces expression of a limited subset of activity-dependent genes, likely resulting from suppressed activity of µOR-expressing GABAergic neurons that limit dopamine neuron firing rate under baseline conditions. In contrast, morphine experience engages largescale transcriptomic alterations in glial populations, including induction of glucocorticoid response genes such as *Fkbp5*. These changes are likely driven by the ability of morphine to increase corticosterone, as glial populations exhibit minimal expression of genes coding for the major opioid receptor classes. These robust glial transcriptional responses highlight the importance of considering non-neuronal cells as active participants in opioid-induced adaptations rather than passive bystanders. Finally, the identification of glucocorticoid-mediated pathways as key drivers of glial responses to opioid exposure reveals a previously underappreciated mechanistic link between stress hormone signaling and the development of addiction-relevant neurobiology. Overall, our findings suggest that targeting glucocorticoid signaling in glial populations may be critical for the development of more comprehensive and effective treatment strategies for opioid use disorder.

## METHODS

### Animals

Male and female 8-12 week old adult Sprague-Dawley rats (16M/16F; ∼200-300g) were purchased from Charles River Laboratories (Wilmington, MA) and pair-housed in an AAALAC-approved animal care facility during the experiment. Rats were maintained on a 12 h light/dark cycle with ad libitum access to food and water and were briefly handled for 3 days prior to beginning baseline Von Frey measurements (1-2min/day) to acclimate them to the experimenters. All experimental procedures were approved by the University of Alabama at Birmingham Institutional Animal Care and Use Committee and were conducted in accordance with the National Institutes of Health Guide for the Care and Use of Laboratory Animals.

### *Drugs* for *in vivo CFA and Morphine experiments*

To assess neuropathic pain response, Complete Freund’s Adjuvant (CFA; Sigma Aldrich, St. Louis, MO) or saline vehicle (100 µl) was injected into the plantar surface of one hind paw of male and female rats (n = 8/sex/treatment group). Mechanical sensitivity was then evaluated 48 hours and 7 days after the injection using the Von Frey assay.

### Von Frey assessment of pain sensitivity

Saline vehicle or CFA (100% i.p.l.) was injected into the hind paw of rats resulting in chronic inflammation as previously described^36^. To monitor changes in pain sensitivity using the Von Frey test, rats were placed in a transparent Plexiglas cubicle with perforated flooring and habituated to the chambers for 20 minutes. Following habituation, nylon monofilaments of increasing weight (Stoelting Touch Test Sensory Evaluator Kit #2 to #9; ∼2.0-60g; Wood Dale, IL) were pressed against the plantar surface of the affected paw and withdrawal threshold was recorded using the up-down method of Dixon^76^.

### Tissue collection from adult rat VTA

One hour after systemic injections of saline or morphine (10mg/kg) rats were euthanized by live decapitation and whole brains were briefly flash frozen in 2-methyl butane. Chilled brains were then blocked into 1-mm-thick coronal sections on a combination of dry and wet ice. Sections containing the VTA (ranging from -4.92-6.46 anterior-posterior (AP) to Bregma) were transferred to glass microscope slides on dry ice, and the VTA was microdissected away from neighboring brain regions (n = 3-4/sex/treatment group; as in^38^. Dissected VTA tissue was then transferred to ice-cold centrifuge tubes, and stored at -80°C until the day of 10X GEM capture.

### Tissue dissociation and GEM capture for snRNA-seq

Frozen microdissected VTA tissue was prepared as previously described^38^. Briefly, tissue was thawed on wet ice and chopped with a scalpel prior to being transferred to 5 ml of ice cold lysis buffer (10 mM tris-HCl, 10 mM NaCl, 3 mM MgCl2 and 0.1% Igepal in nuclease-free water (Sigma-Aldrich, 18896-50ML). After 15 min the lysis was quenched with 5 ml of complete Hibernate A media (Thermo Fisher Scientific, A1247501) supplemented with B27, GlutaMAX (Life Technologies, 35050-061), and NxGen RNase Inhibitor (0.2 U/m; Lucigen, 30281-2). Tissue was then triturated by fire-polished Pasteur pipette (three pipettes of decreasing diameter) and filtered with a 40µm pre-wet cell strainer. Samples were pelleted and then washed in nuclei wash and resuspension buffer (phosphate-buffered saline (PBS), 1% bovine serum albumin (BSA), and NxGen RNase Inhibitor). The resulting pellet was resuspended in 800 ml of wash buffer and 7-aminoactinomycin D (7-AAD; Thermo Fisher Scientific, 00-6993-50) was added prior to fluorescence-activated cell sorting (FACS) on a FACSAria instrument (70 µm nozzle, BD Biosciences). Immediately after FACS, nuclei were washed a final time at 250 rcf in 10 ml of supplemented Hibernate A containing 1% BSA and RNase Inhibitor for 5 min at 4C. Nuclei were brought to a concentration of 1,000 nuclei/µL. A total of 8,000 nuclei pooled from two rats (4,000 male nuclei/4,000 female nuclei from the same treatment group) and were loaded into individual wells of the Chromium NextGem Single Cell Chip (10x Genomics, catalog no. 10000121).

### 10x GEX library preparation and sequencing

Libraries were constructed according to manufacturer’s instructions for Chromium Next GEM single cell 3’ library and gel bead kit (10x Genomics, v3.1 single index, catalog no. 10000121, version 3 chemistry). A total of 50,513 nuclei were captured across 16 GEM (gel bead in emulsion) wells in total (4 wells per treatment group) using the 10x Chromium Controller. Each GEM well contained nuclei from 1 male and 1 female animal combined by treatment group for a total of 16 rats per sex. Libraries were sequenced on an Illumina NovaSeq6000 (S2 flow cell) to an average read depth of ∼71,000 read pairs per nucleus (range 48,000 - 123,000).

### snRNA-seq analysis

Raw sequencing data for each GEM well was processed and aligned to the rat genome (Rn7) in Cell Ranger^77^ v.9.0.1 on the Cheaha high performance computing cluster at the University of Alabama at Birmingham, with the associated Ensembl gene transfer format file (version 105) modified to add annotation for the *Xist* gene. CellRanger filtered outputs were analyzed with Seurat^78^ v.5.2.1 in R^79^ v.4.4.0. Ambient RNA was estimated and removed using SoupX^80^ v.1.6.2. Cells with <200 genes and >5% of reads mapping to the mitochondrial genome were removed. For each GEM well, counts were normalized, scaled by a factor of 10,000, and log-transformed. Doublets were identified and removed using scDblFinder^81^ v.1.18.0 with the default expected doublet rate of 1% per 1,000 cells. To predict the sex of cells from pooled GEM wells, a logistic regression model was fit and validated using previously published rat VTA snRNA-seq data^82^ with the glm() function of the stats package (v.4.2.0) in R. Cells were classified as male if model predicted probabilities were < 0.25, and female if > 0.75. Low-confidence cell sex classifications were labeled as unknown if predicted probabilities were between 0.25-0.75 and omitted from downstream analysis. We then integrated all samples in Harmony^83^ v.1.2.3, using principle component analysis PCA) cell embeddings with 50 principle components (PCs). Downstream analysis was performed on the integrated object. For each gene, log-normalized counts were scaled to a mean of 0 and variance of 1 across cells. Dimensionality reduction was conducted via PCA retaining 20 PCs, followed by UMAP with the same 20 PCs. Cells were then clustered following the Seurat workflow, generating a K-nearest neighbor graph and clustering under a Louvain algorithm with a resolution of 0.4. These parameters were chosen by sweeping a range of PC (10-25) and resolution (0.1-0.5) values and selecting the one that best recapitulated known VTA cell types. Local Inverse Simpson’s Index (LISI) scores were computed to assess sample integration across clusters using lisi v1.0 in R^83^. To refine resolution of VTA neuronal subtypes and assign cells to classes, we followed the Allen Institute’s Whole Mouse Brain taxonomy using the MapMyCells web interface^84^ (RRID:SCR_024672). After removing additional low-quality or ambiguous cell populations, we repeated the dimensionality reduction and clustering steps with the same parameters. Cells were considered low quality or ambiguous if: 1) present in small clusters mapping to multiple cell classes in MapMyCells with low UMI/gene counts and elevated mitochondrial reads, 2) flagged as low-confidence oligodendrocytes in MapMyCells and clustered outside of oligodendrocyte populations, or 3) were assigned to clusters containing <100 cells, which were considered low-confidence due to insufficient representation. Differential gene expression analysis was conducted in DESeq2 v.1.44.0 to test for effects of treatments (CFA, morphine) and treatment interaction, with sex as a covariate. We generated a pseudobulked counts matrix summing counts across all cells per sample in each cell type. Genes with < 5 total counts were discarded from testing. To identify significantly enriched GO terms and KEGG and REACTOME pathways in the sets of glial DEGs, we conducted enrichment analyses in R using clusterProfiler^85^ v.4.12.6 (for GO and KEGG) and ReactomePA^86^ v.1.48.0 (for REACTOME) with the rat genome. We separately tested up- and downregulated morphine DEGs for astrocytes, microglia, and the Olig.1 cluster, as well as the concatenated set of DEGs for those three glial clusters. We also tested the treatment interaction DEGs for astrocytes. For all analyses, the background gene set consisted of all detected genes in the full Seurat object.

### C6 cell culture

C6 cells used for in vitro rat glia experiments were acquired from American Type Culture Collection (ATCC; cat # CCL-107) and maintained in Ham’s F12k-based medium (Thermo Fisher Scientific, cat # 21127022) supplemented with 2.5% fetal bovine serum and 12% horse serum. Cells were seeded at a density of 95,000 cells/well in 12-well plates and allowed to reach 70-90% confluency over 2 days. Pharmacological treatments (e.g., corticosterone, mifepristone, DAMGO) were then delivered in triplicate (3 wells/treatment group, across 2 culture experiments). After treatments, media was aspirated from the cells and plates were frozen at -80 °C until RNA extraction for downstream RT-qPCR analysis.

### Human genetically immortalized neural progenitor cells (NPCs)

The human NPC line ReNcell VM (Millipore Sigma, cat. # SCC008; RRID: CVCL_ E921) was originally derived from the ventral mesencephalon of a 10-weeks gestation euploid (46, XY) fetus, in accordance with applicable ethical and legal guidelines, and immortalized by retroviral transduction with the v-myc oncogene^87^. Ploidy and chromosomal sex were confirmed by karyotype. Cells were grown on cell culture plates coated with laminin (10 μg/mL; Millipore Sigma, cat. # L2020) in ReNcell NSC Maintenance Media (Millipore Sigma, cat. # SCM005) supplemented with recombinant human epidermal growth factor (EGF, 20 ng/mL; STEMCELL Technologies, cat. # 78006), recombinant human basic fibroblast growth factor (bFGF, 20 ng/ mL; STEMCELL Technologies, cat. # 78003), and penicillin-streptomycin (Fisher Scientific, cat. # 15-140-122). Cells were passaged every 3-4 d at 85-90% confluency using StemPro Accutase Cell Dissociation Reagent (Gibco, cat. # A1110501) and were used for no more than 10 passages.

### Differentiation of human NPCs into astrocytes

Differentiation of NPCs into astrocytes was adapted from previously published protocols^88^. NPCs were seeded onto cell culture plates coated with poly-L-lysine (50 μg/mL; Millipore Sigma, cat. # P4707) supplemented with laminin (20 μg/mL) and grown overnight in ReNcell NSC Maintenance Media supplemented with EGF, bFGF, and the Rho kinase (ROCK) inhibitor Y-27632 (10 μM; STEMCELL Technologies, cat. # 72304). Astrocyte differentiation commenced the next day (DIV1) by performing a full media change into DMEM/F-12 (Thermo Fisher Scientific, cat. # 11320033) supplemented with N-2 Supplement (1X) (Thermo Fisher Scientific, cat. # 17502048), GlutaMAX (1X) (Thermo Fisher Scientific, cat. # 35050061), recombinant human ciliary neurotrophic factor (CNTF; 20 ng/mL) (STEMCELL Technologies, cat. # 78010), recombinant human bone morphogenetic protein 4 (BMP4, 10 ng/mL; PeproTech, cat. # 120-05), and penicillin-streptomycin. Beginning on DIV3, half-media changes were performed every 2-3 d until DIV14, with visual microscopic inspection preceding each media change to monitor cell survival and morphological maturation.

### In vitro glucocorticoid and opioid pharmacology

To engage GR signaling *in vitro*, human-derived astrocyte cultures were treated on DIV14 with cortisol (5 μM; Sigma-Aldrich, cat. # C-106) or dexamethasone (100 nM; Sigma-Aldrich, cat. # D4902) for 5 h. Vehicle-treated (methanol) groups were included as negative controls. Where indicated, cells were pretreated for 30 min with the GR antagonist mifepristone (also known as RU-486, 1 μM; Sigma-Aldrich, cat. # 475838) or vehicle (ethanol). The relative contributions of opioid *versus* glucocorticoid signaling were assessed using the selective μ-opioid receptor agonist DAMGO ([D-Ala^2^, *N*-Me-Phe^4^, Gly^5^-ol]-enkephalin; Sigma-Aldrich, cat. # E7384) at concentrations ranging from 100 nM to 10 μM in tenfold increments. Following treatments, media was aspirated and cells were immediately stored at -80°C or subjected to RNA extraction for downstream gene expression profiling.

### CRISPR-dCas9 and sgRNA construct design

For transcriptional repression experiments using CRISPR inhibition (CRISPRi), a lentivirus plasmid previously optimized for robust neuronal expression^89^ was modified in-house using restriction digest cloning to replace the human synapsin promoter (hSYN) with astrocyte-selective promoter GfaABC1D^75^. Gene-specific sgRNA targets were designed using online tools provided by CHOPCHOP (http://chopchop.cbu.uib.no/). To ensure specificity, CRISPR RNA (crRNA) sequences were analyzed with National Center for Biotechnology Information’s Basic Local Alignment Search Tool. Briefly, crRNAs were annealed and ligated into the Sp dCas9 sgRNA scaffold using BbsI cut sites. Cloned plasmid products were sequence-verified with Oxford Nanopore long-read sequencing (Plasmidsaurus). On DIV10, human-derived astrocytes were co-transduced overnight with lentiviruses expressing GfaABC1D-dCas9-KRAB-MeCP2 and sgRNAs targeting either the bacterial gene lacZ as a negative control or human *NR3C1*. On DIV 11 and 12, cells were imaged to verify virus expression before moving forward with cortisol treatments and/ or RNA extractions on DIV14.

### Lentivirus production

Lentiviruses were produced as described previously (Savell et al., 2019) in a sterile environment subject to BSL2 safety conditions by co-transfecting human embryonic kidney 293T cells (ATCC, cat. # CRL-3216) with the specified CRISPR plasmid, the psPAX2 packaging plasmid, and the pCMV-VSV-G envelope plasmid (Addgene plasmids12260 and 8454) with FuGENE HD (Promega) for 40 to 48 hours in a 12-well culture plate. Supernatant containing viral particles was concentrated using Lenti-X concentrator (Takara), resuspended in sterile PBS, and used immediately or stored at -80°C until use.

### Immunocytochemistry

For validation of successful differentiation of human NPCs into astrocytes we performed immunocytochemistry as described previously^90^ for NPC markers (NESTIN and SOX2) and astrocyte markers (GFAP and S100B) at DIV 1 and DIV10 in the NPC differentiation protocol. Similarly, to validate expression of CRISPRi machinery in astrocytes, we performed immunocytochemistry for FLAG and GFAP at DIV14. Media was aspirated off cells and wells were washed briefly with 1x PBS (pH 7.4). Cells were fixed with 4% paraformaldehyde (PFA; pH = 7) in PBS at room temperature for 25 minutes on a rocking platform. Ice cold PBS was used to wash cells twice following PFA. PBS with .25% Triton X-100 was added to cells for 20 minutes to permeabilize membranes. Cells were washed thrice with PBS for 5 minutes and then incubated in a blocking buffer (PBS with 10% Thermo Blocker BSA, 1% goat serum, 0.05% Tween-20, and 300 mM glycine) for 60 min. Next, cells were incubated in primary antibody buffer with anti-NESTIN antibody (1:2400 in PBS with 10% Thermo Blocker BSA and 1% goat serum, 33475S, Cell Signaling Technology), anti-SOX2 antibody (1:400 in PBS with 10% Thermo Blocker BSA and 1% goat serum, 3579S, Cell Signaling Technology, RRID AB_2195767), anti-S100B antibody (1:200 in PBS with 10% Thermo Blocker BSA and 1% goat serum, EP1576Y, Abcam, RRID AB_882426), anti-GFAP antibody (1:1000 in PBS with 10% Thermo Blocker BSA and 1% goat serum, PA1-10019, Thermo Fisher Scientific, RRID AB_1074611), and/or anti-FLAG antibody (1:250 in PBS with 10% Thermo Blocker BSA and 1% goat serum, MA1-91878, Thermo Fisher Scientific, RRID AB_1957945) overnight at 4°C, with rocking. Plates sealed in parafilm and wrapped in foil to prevent photobleaching. The next day, the cells are washed with PBS with .25% TritonX-100 for 10 min, and they were rinsed twice with PBS for 5 minutes afterwards. The secondary antibody buffers with Goat anti-Mouse Alexa Fluor 488 (1:1000 in PBS with 10% Thermo Blocker BSA and 1% goat serum, A-10667, Thermo Fisher Scientific, RRID AB_2534057) and/ or Goat anti-Rabbit Alexa Fluor 546 (1:500, A-11010, Thermo Fisher Scientific, RRID AB_2534077) were added to the wells and allowed to incubate for 1 hour at room temperature under foil to protect from light. Following incubation, wells were washed three times with PBS. Glass coverslips were added to wells along with Prolong Diamond Antifade Mountant with DAPI (P36962, Thermo Fisher Scientific) to stain nuclei. Images were taken on a Nikon TiS inverted fluorescence microscope at 20x (Fig. 5) or 10x (Fig.6) magnification.

### RNA extraction and RT-qPCR

RNA was extracted from rat C6 or human-derived astrocytes per manufacturer’s instructions using the Qiagen RNeasy RNA extraction kit (#74106, Qiagen, Hilden, Germany), and reverse transcription into cDNA was performed using iScript Supermix (Bio-Rad #1708840). Expression of rat *Gapdh* and *Fkbp5* or human *GAPDH, FKBP5*, and *NR3C1* was measured using quantitative PCR amplification in duplicate reactions with SsoAdvanced Universal SYBR Green Supermix (BioRad) on a CFX96 real-time PCR system (Bio-Rad) at 95°C for 3min, followed by 40 cycles of 95°C for 10 s and 58°C for 30 s, followed by real-time melt analysis to verify product specificity. Quantification was performed as previously described^91^. Primer sequences are listed below.

Rat *Gapdh* Forward:

ACCTTTCATGCTGGGGCTGGC

Rat *Gapdh* Reverse:

GGGCTGAGTTGGGATGGGGAC

Rat *Fkbp5* Forward:

TGGTCTGACTCTCGTGTTTCTTG

Rat *Fkbp5* Reverse:

CGCAGGGTGTACGCCAAC

Human *GAPDH* Forward:

TGTCAAGCTCATTTCCTGGTAT

Human *GAPDH* Reverse:

CTCTCTTCCTCTTGTGCTCTTG

Human *FKBP5* Forward:

CTCCATGTGCCAGAAAAAGGC

Human *FKBP5* Reverse:

CCGAGTTCACTGGGACTCTTC

Human *NR3C1* Forward:

GGAATAGGTGCCAAGGATCTGG

Human *NR3C1* Reverse:

GCTTACATCTGGTCTCATGCTGG

## Statistical analyses

Statistical analyses were completed and graphed using GraphPad Prism software. Behavioral effects of CFA-induced inflammatory pain were analyzed using a two-way analysis of variance (ANOVA) with Tukey’s post hoc comparisons. Differences in IEG expression between neuronal subpopulations in Fig. 2e were calculated using one sample t-tests. Gene expression differences from RT-qPCR experiments were compared with unpaired t-tests or one-way ANOVAs with Tukey’s post hoc comparisons where appropriate.

## DATA AVAILABILITY

All data needed to evaluate the conclusions in the paper are present in the paper and/or the Supplementary Materials. Custom code can be found at https://github.com/Jeremy-Day-Lab/Tuscher_etal_VTA_Pain_Morphine. Sequencing data that support the findings of this study are available in Gene Expression Omnibus (GSE308304). Single nucleus RNA sequencing data from this experiment can also be freely viewed at www.ratlas.org, and the Seurat R object for this dataset can be found at https://doi.org/10.5281/zenodo.17138061.

## ACKNOWLEDGEMENTS

We thank all Day Lab members for assistance and support. We acknowledge support from the University of Alabama at Birmingham Biological Data Science Core (RRID:SCR_021766), the UAB Heflin Center for Genomic Sciences, and the UAB Flow Cytometry & Single Cell Core Facility.

## AUTHOR CONTRIBUTIONS

Conceptualization: JJT, RAP, RES, JJD

Methodology: JJT, RAP, NJR, RES, JJD

Investigation: JJT, AC, RAP, NJR, RES, JJD

Formal analysis: JJT, AC, RAP, CEN, GT, LI, RES, JJD

Software: LI

Resources: RES, JJD

Funding acquisition: RES, JJD

Data curation: JJT, CEN, RAP, GT, JJD

Visualization: JJT, AC, RAP, CEN, GT, JJD

Validation: JJT, RES, JJD

Project administration: JJT, RES, JJD

Supervision: JJT, RES, JJD

Writing—original draft: JJT, RES, JJD

Writing—review & editing: JJT, AC, RAP, CEN, GT, NJR, LI, RES, JJD

---

**Figure S1.**
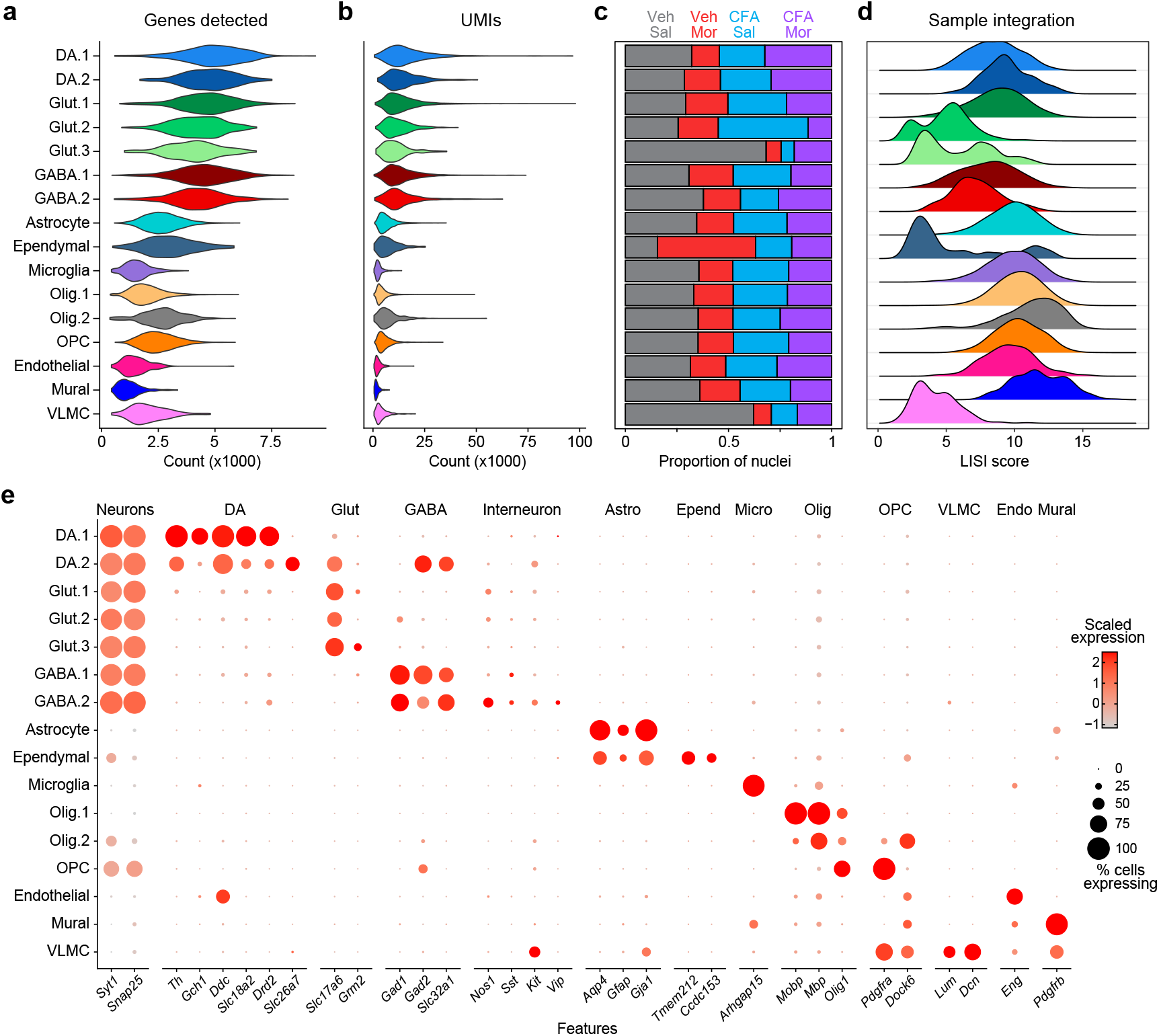
Quality control, integration, and marker gene analysis for snRNA-seq dataset. **a**, Violin plot showing number of genes detected in each cluster ID. **b**, Violin plot of the number of unique molecular identifiers (UMIs) in each cluster ID. **c**, Bar graph of the proportion of nuclei contributed from each experimental group. **d**, Local Simpson’s Inverse Index (LISI) scores for each annotated cluster. **e**, Dot plot of expression level and percentage of known marker genes in each cluster.

**Figure S2.**
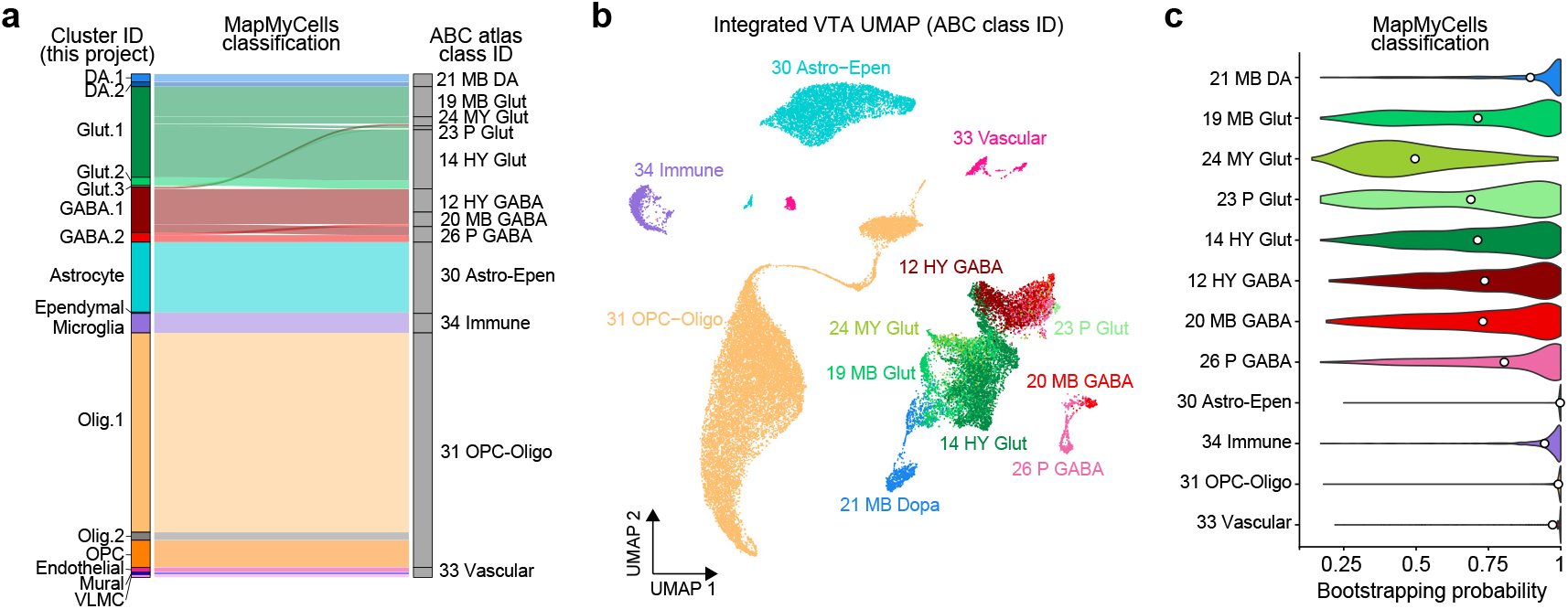
Comparison of annotated cluster IDs with the Allen Brain Cell (ABC) mouse brain atlas. **a**, Alluvial plot showing relationship between cluster ID assignments in this project and ABC atlas class ID. **b**, Integrated VTA UMAP showing predicted Allen class ID for each nucleus. **c**, Bootstrapping probability for predictions for each ABC class ID indicates lower confidence in predictions for glutamatergic and GABAergic neurons.

**Figure S3.**
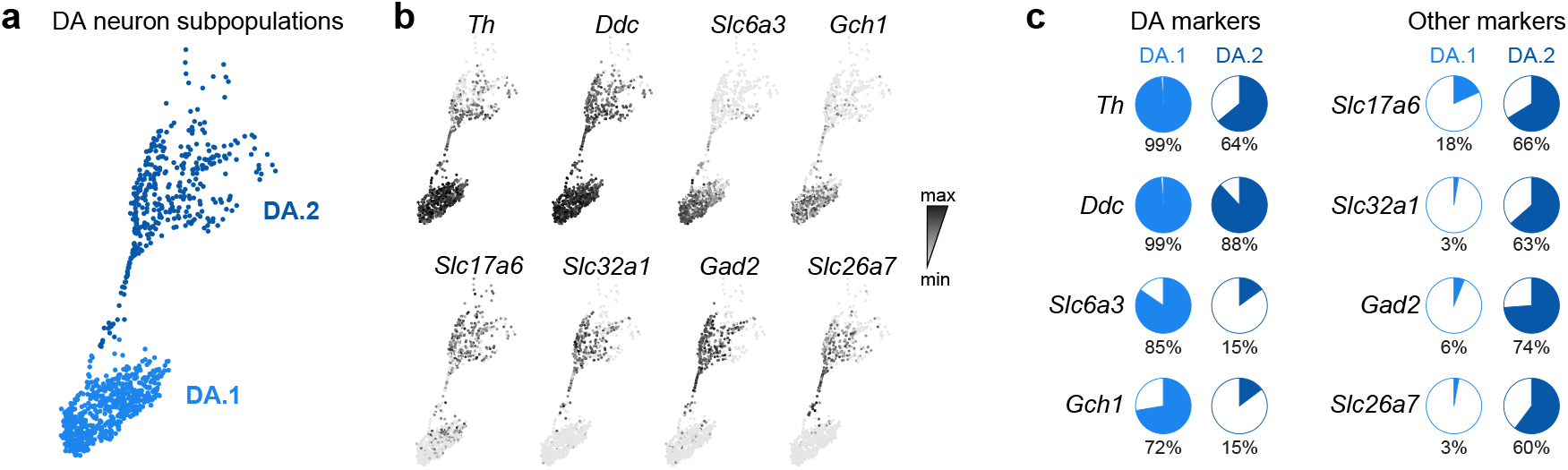
Dopamine neuron subtype marker comparison. **a**, Feature plot of dopamine neuron populations with cluster IDs. **b**, expression of dopamine marker genes (top row) and glutamate and GABA marker genes (bottom row). **c**, Pie graphs showing percent of DA.1 or DA.2 neurons expressing DA and other marker genes. DA.1 neurons express high levels of *Gch1* and little *Slc26a7*, whereas DA.2 neurons express high levels of *Slc26a7* and little *Gch1*.

**Figure S4.**
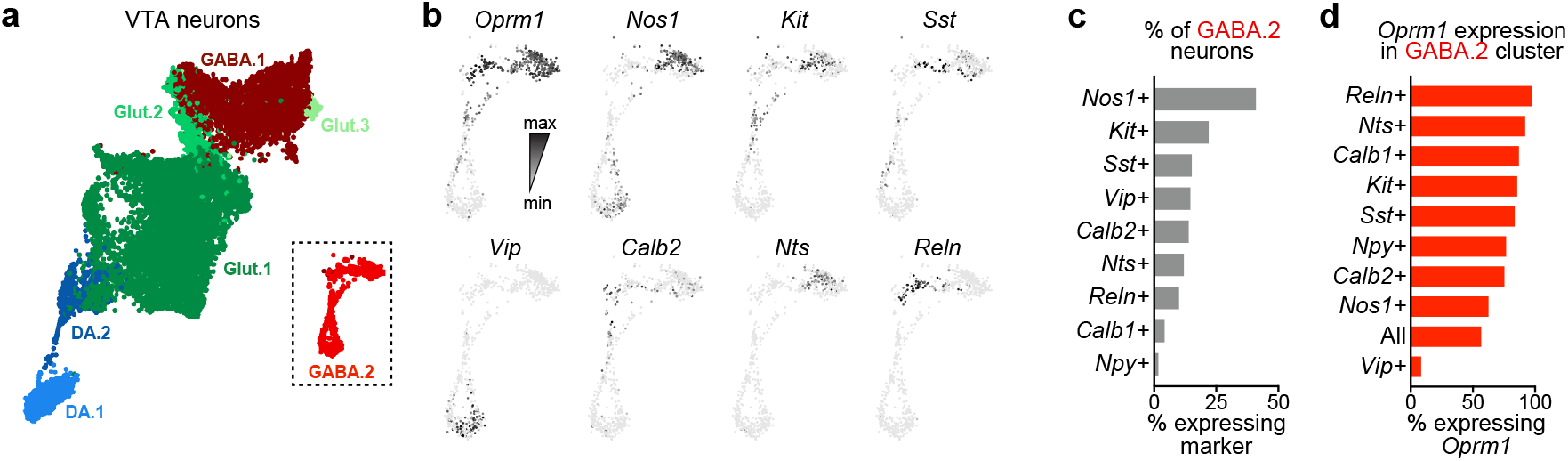
Overlap of GABAergic interneuron markers with *Oprm1* expression in the VTA GABA.2 cluster. **a**, UMAP of VTA neuronal populations. **b**, Expression feature plots of *Oprm1* and known interneuron markers within the GABA.2 cluster. **c**, Percentage of GABA.2 neuronal population expressing common interneuron markers. **d**, Percentage of neurons with interneuron marker expression that also express *Oprm1*.

**Figure S5.**
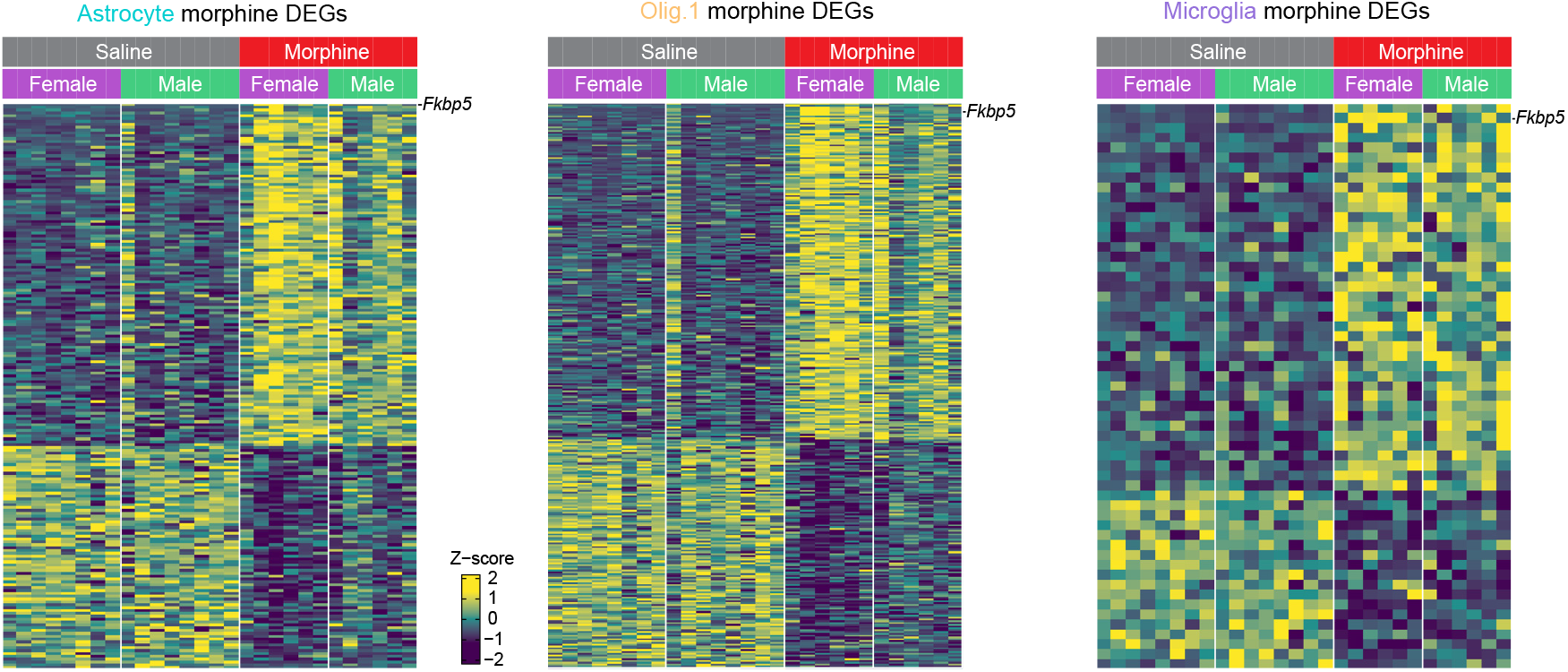
Heatmaps of DEGs in Astrocyte, Olig.1, and Microglia clusters with individual sample level data shown and split by sex. Each row represents a single DEG, and columns represent individual samples. For each gene, pseudobulked log-normalized counts were centered and scaled across samples to calculate the Z-score using the formula Z = (x - μ) / σ, where μ is mean and σ is standard deviation.

**Figure S6.**
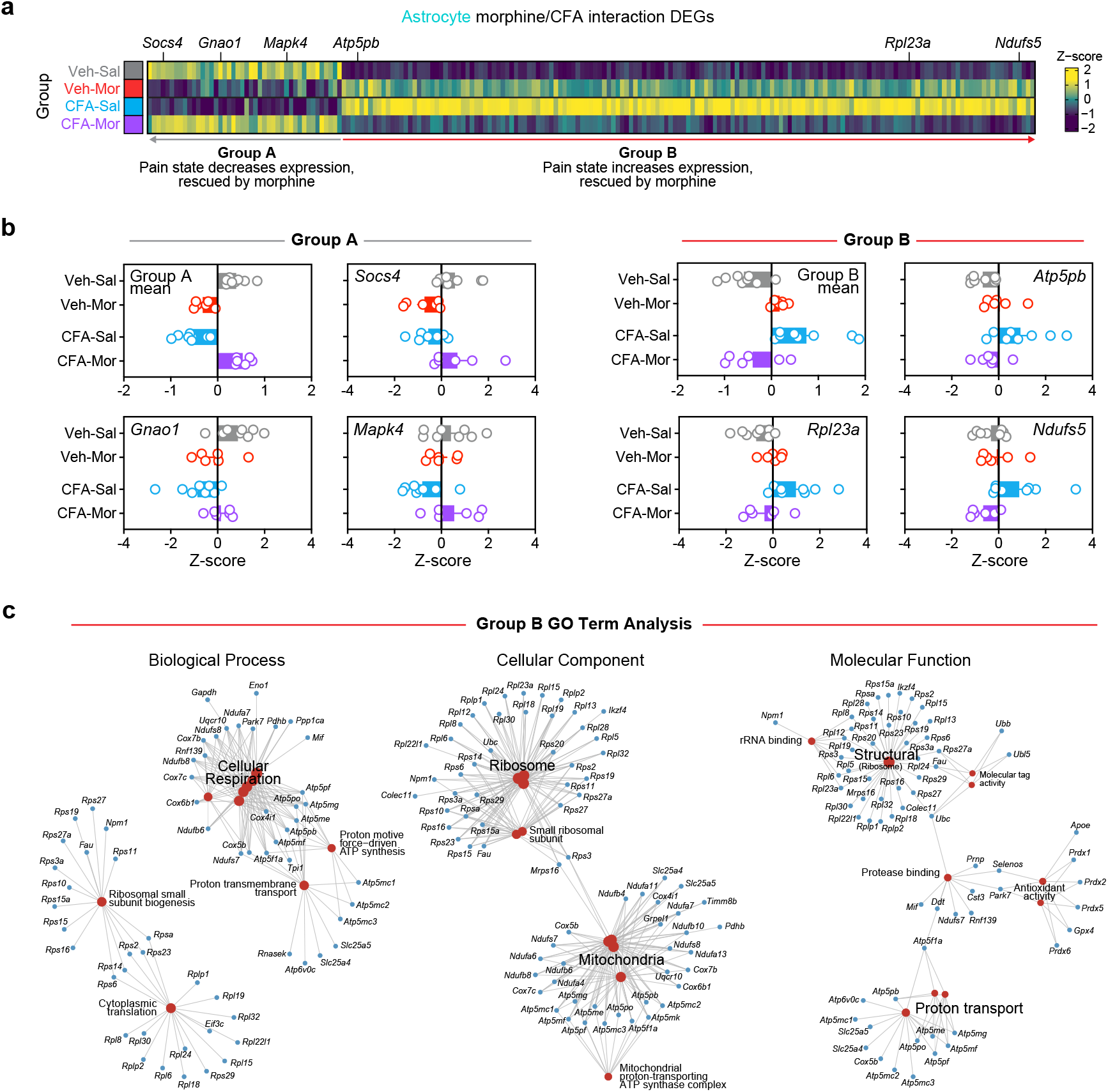
Pain and morphine interact to alter gene expression in VTA astrocytes. **a**, Heatmap of Astrocyte cluster CFA x morphine interaction DEGs. Each column represents a single DEG, and rows represent group average values. For each gene, pseudobulked log-normalized counts were centered and scaled across groups to calculate the Z-score using the formula Z = (x - μ) / σ, where μ is mean and σ is standard deviation. **b**, Mean and representative interaction DEGs belonging to Group A and Group B. **c**, Network diagram showing enriched gene ontology terms in Group B interaction DEGs.

